# Pelvic spine reduction affects diet but not gill raker morphology in two polymorphic brook stickleback (*Culaea inconstans*) populations

**DOI:** 10.1101/2023.04.03.535428

**Authors:** Jonathan A. Mee, Emily Yap, Daniel M. Wuitchik

**Affiliations:** Department of Biology, Mount Royal University, 4848 Mount Royal Gate, Calgary, Alberta, T3E 6K6, Canada; Department of Biology, Boston University, 5 Cummington Mall, Boston, Massachusetts, 02215, United States

## Abstract

Pelvic spine polymorphism occurs in several species in the stickleback family (Gasterosteidae). Given the similar phenotypic polymorphisms in multiple stickleback species, we sought to determine the extent of parallelism in the ecological correlates of pelvic spine reduction. Based on a metabarcoding analysis of brook stickleback gut contents in two polymorphic populations, we found significant diet differences were associated with pelvic spine reduction, but we found no clear or consistent trend supporting a tendency for benthic feeding in pelvic-reduced brook stickleback. These results contrast with those found in threespine stickleback where pelvic spine reduction is often associated with a benthic diet. Hence, we found non-parallel consequences of spine polymorphism across species. Furthermore, a difference in gill raker morphology has been frequently observed between ecomorphs with differen diets in many fish species. But, we found no evidence of any difference in gill raker morphology associated with pelvic spine polymorphism in brook stickleback.

## Introduction

The maintenance of polymorphisms in natural populations is a central concept in evolutionary biology. Polymorphisms are of particular interest when they are maintained at a frequency that is too high to be explained by mutation-selection balance (Ford 1945, Lande 1975). In cases where a polymorphism persists in a population at high and stable frequency, the maintenance of the polymorphism is typically explained by some form of balancing selection or by heterosis (Dobzhansky 1971, Takahata et al. 1992, Rainey et al. 2000, Gigord et al. 2001, Punzalan et al. 2005). These cases are of interest because they afford the opportunity to study the genetic and ecological mechanisms that result in the balance of selective forces that maintain diversity. For example, balancing selection, in the form of pollinator-mediated negative frequency dependent selection, maintains a stable polymorphism in flower colour in the rewardless orchid *Dactylorhiza sambucina* (Gigord et al. 2001). Regardless of whether or not a given polymorphism is maintained by balancing selection (e.g. by frequency-dependent selection or temporally-fluctuating selection), a polymorphism has the potential to affect other traits via pleiotropy, and a polymorphism may cause divergence in a suite of traits via plastic or evolutionary responses to the ecological or behavioural consequences of the polymorphism.

Studies of polymorphisms in threespine sticklebacks have provided foundational contributions to our understanding of evolutionary processes (Schluter and McPhail 1992, Reimchen 1994, Schluter 2000, Bolnick 2004, Colosimo et al. 2005, Chan et al. 2010, Hendry et al. 2013, Schluter et al. 2022). The establishment of threespine stickleback as a model system in evolutionary biology (Reid et al. 2021) has also increased interest in the study of adaptive polymorphisms within other species in the stickleback family, Gasterosteidae (Blouw and Boyd 1992, Shapiro et al. 2009, Klepaker et al. 2013, Shikano et al. 2013, Lowey et al. 2020, Jeffries et al. 2022, Willerth et al. 2022). For example, spine reduction or loss has evolved independently in four stickleback genera that share a common ancestor approximately 27 million years ago: *Gasterosteus*, *Pungitius*, *Apeltes*, and *Culaea* (Nelson 1969, Nelson 1977, Hagen and Blouw 1983, Blouw and Hagen 1984a, Chan *et al*. 2010, Varadharajan et al. 2019). The involvement of predation by birds, fish, mammals, and insects as a selective agent on spines has been established by many studies involving multiple stickleback species (Hoogland et al. 1957, Hall 1956, Reisman and Cade 1967, Reimchen 1980, Reist 1980*a*, Reist 1980*b*, Reist 1983, Blouw and Hagen 1984b, Foster et al. 1988, Reimchen 1992, Reimchen 1994, Vamosi 2002, Rundle et al. 2003, Vamosi and Schluter 2004, Marchinko 2009, Lescak and von Hippel 2011, Lescak et al. 2012, Miller et al. 2017). These studies typically take advantage of the existence of variation among individuals within a population (or within sympatric or parapatric populations) to determine which pelvic phenotype (e.g. spined or unspined) is favoured by a given predator. For example, several studies have shown that gape-limited predators, such as trout (*Oncorhynchus mykiss*) or juvenile pike (*Esox lucius*), select for increased spine length or against spine loss, whereas predators that are not gape-limited, such as adult pike or larvae of *Dytiscus* spp. (water beetles), select for reduced spines or spine loss (Reist 1980a, 1980b). Given that multiple predators occur in an environment (e.g. a lake may contain insects, fish, and birds that prey on stickleback; see Reimchen 1994), and that predators are not uniformly distributed within an environment across space, seasonsm, or years (Reimchen 1994), spine polymorphism within stickleback populations may be maintained by a balance of selection driven by multiple predators (e.g. gape-limited and non-gape-limited) across multiple habitats or multiple points in time. There is also evidence that spines may be selected against in calcium-deficient environments, presumably due to the resource cost of gowing these bony structures (Giles 1983, Bell et al. 1993, Spence et al. 2013, Haines et al. 2023). So, spine polymorphism may be maintained by a balance of selection for spines by gape-limited predators and against spines due to physiological costs in low-calcium environments (e.g. in some freshwater lakes).

The ecological correlates of spine polymorphism have been well-studied and characterized in threespine stickleback (Bell and Foster 1994, Klepaker et al. 2013, Haines et al. 2023). These ecological correlates include differences between spine morphs in predation, habitat preference, and diet, which, collectively or invidually, give the opportunity for additional adaptive or plastic morphological or behavioural differences between spine morphs. Pelvic spine divergence in some lakes is known to be associated with morphological and ecological divergence between reproductively isolated limnetic and benthic threespine stickleback, wherein the benthic species often evolves reduced spines (Schluter and McPhail 1992, Hatfield and Schluter 1999, Schluter 1993, Hatfield 1997, Boughman 2001, Peichel et al. 2001, Rundle and Schluter 2004). Even without reproductive isolation, a well-established difference associated with spine reduction in threespine stickleback is that spined stickleback consume more planktonic food sources and unspined stickleback consume more benthic food items, which is associated with a difference in gill raker morphology where fewer, shorter, and thicker gill rakers are found in benthic feeding fish (Reimchen 1980, McPhail 1984, Ridgeway and McPhail 1984, Schluter 1993, Nagel and Schluter 1998, Rundle et al. 2003, Matthews et al. 2010). It is unclear, however, whether the reduction in spines that has occurred in other stickleback species is associated with parallel ecological and morphological changes, such as parallel changes in diet or gill raker morphology.

In brook stickleback (*Culaea inconstans* Kirtland, 1840), within-population pelvic spine polymorphism does not involve reproductively isolated morphs (Lowey et al. 2020) and the ecological and morphological changes associated with pelvic spine reduction are not necessarily consistent among populations (Willerth et al. 2022). To date, however, no study has directly addressed whether pelvic spine reduction is associated with a difference in diet in brook stickleback in a parallel manner to the difference in diet in pelvic-reduced relative to spined threespine stickleback. Hence, the extent of ecological or phenotypic parallelism among stickleback species associated with the reduction or loss of pelvic spines is unknown. Resolving this question about the extent of parallelism among stickleback species will help address more general questions about the extent of phenotypic parallelism across taxa (Bolnick et al. 2018, De Lisle and Bolnick 2020).

In this study, we sought to understand whether pelvic spine reduction is correlated with ecological and morphological changes in brook stickleback. We addressed the hypothesis that pelvic spine polymorphism is associated with a difference in diet in brook stickleback, and we predicted that, as in threespine stickleback, pelvic-reduced fish eat a more benthic diet. To address our hypothesis, we used metabarcoding of gut contents to compare diet among spined and pelvic-reduced brook stickleback in two polymorphic populations. Because gill raker morphology differs between ecomorphs of threespine stickleback and many other fish species (see, for example, Mee et al. 2015), we also hypothesized that gill raker morphology would differ between spined and pelvic-reduced brook stickleback.

## Methods

### Sample collection and study site

Brook stickleback were collected in Alberta, Canada, from Muir Lake (UTF-8 encoded WGS84 latitude and longitude: 53.627659, -113.957524) and Shunda Lake (52.453899, - 116.146192) in the summer of 2017 and 2019 using unbaited minnow traps. To sample fish from the littoral zone, where they were more likely to have a benthic diet, traps were set within throwing distance from the shore (maximum 4m to 5m from shore) at a depth of 0.5m to 2m. To sample fish from the limnetic zone, where they were more likely to have a planktonic diet, traps were set at least 50m from shore and suspended from floats at a depth of 1-2m. All traps were retrieved within one to twelve hours after being set. In each year, all brook stickleback were retained until we had collected 40 (in 2017) or 30 (in 2019) spined individuals. Then, once we had captured 30 or 40 spined fish, we retained only pelvic-reduced individuals until we had achieved a balanced sample of spined and pelvic-reduced morphotypes (Table 1). Pelvic morphology was assessed at the time of capture via close inspection of the pelvic region of each fish with forceps, and sex was identified at the time of capture based on the presence of male nuptial colouration or the presence of eggs. Each fish included in this study was euthanized by immersion in an overdose mixture of lake water and eugenol. In 2019, the entire gut of each fish was removed and preserved in 95% ethanol. The body of each fish was preserved in 70% ethanol. As described in Willerth et al. (2022), we stained each fish with Alizarin red to observe calcium-containing osteocytes more clearly (such as in pelvic bones, spines, branchial arches, and gill rakers) and we confirmed pelvic morphology in the lab using magnified digital images.

**Table 1.** Summary of samples analyzed in this study. Numbers in parentheses indicate the samples used in the gut metabarcoding analysis.

|  |  | Pelvic Phenotype |  |  |
| --- | --- | --- | --- | --- |
|  |  | Spined | Vestigial | Unspined |
| 2017 |  |  |  |  |
| Muir Lake |  |  |  |  |
|  | female | 19 | 1 | 25 |
|  | male | 22 | 1 | 18 |
| Shunda Lake |  |  |  |  |
|  | female | 20 | 22 | 6 |
|  | male | 25 | 8 | 3 |
| 2019 |  |  |  |  |
| Muir Lake |  |  |  |  |
|  | female | 3 (3) | 0 | 7 (7) |
|  | male | 25 (18) | 1 | 27 (18) |
| Shunda Lake |  |  |  |  |
|  | female | 13 (13) | 12 (10) | 3 (3) |
|  | male | 17 (12) | 10 (9) | 2 (2) |

The fish collected from Muir Lake and Shunda Lake were used in previous comparisons of stable isotope signatures and morphology between spined and unspined brook stickleback (Willerth et al. 2022). These lakes are well-suited for studies of brook stickleback pelvic spine polymorphism because they contain relatively high abundances of spined and pelvic-reduced morphs (Nelson 1977, Lowey et al. 2020). The pelvic-reduced morphs include unspined individuals that completely lack spines or any vestige of a pelvic girdle, but also individuals with vestigial (a.k.a. intermediate) pelvic structures that lack some pelvic element (e.g. one spine missing, half the pelvic girdle missing, etc.; see Klapaker et al. 2013 for a detailed description of variation in pelvic morphology across stickleback species). In Muir Lake, approximately 40% of the brook stickleback are unspined, with very few (less than 2%) intermediate morphs (Lowey et al. 2020, and personal observation). In Shunda Lake, approximately 20% of the brook stickleback have some sort of pelvic reduction (unspined or intermediate morphs), and approximately 25% of these pelvic-reduced morphs are completely unspined (Lowey et al. 2020, and personal observation). Recent surveys and stocking records (Alberta Environment and Parks 2021) suggest that salmonids, such as brown trout (*Salmo trutta* Linnaeus 1758) and rainbow trout [*Ocorhychus mykiss* (Walbaum 1792)], are present in both lakes, although we also observed other predators, such as loons [*Gavia immer* (Brunnich 1764)], dragonfly nymphs (Gomphidae and other unidentified families), giant water bugs [*Lethocerus americanus* (Leidy 1847)], and backswimmers (Notonectidae) in both lakes (Willerth et al. 2022). Both lakes contain abundant Daphniid zooplankton, and littoral macroinvertebrate sampling revealed Trichoptera larvae, amphipods, leeches, and Chironomidae larvae (Willerth et al. 2022). Sampling permits were issued by the Government of Alberta, and fish handling protocols were approved by the Animal Care Committee at Mount Royal University (Animal Care Protocol ID 101029 and 101795).

### Diet analysis

Fish guts collected from 95 fish in 2019 (Table 1) were shipped to the Canadian Centre for DNA Barcoding at the University of Guelph for membrane-based DNA extraction and amplicon sequencing (Ivanova et al. 2006). This metabarcoding approach used five PCR primer sets designed to amplify a barcode region of the cytochrome c oxidase subunit I (COI) gene in arthropods, mollusks, annelids, amphipods, and microalgae (details of COI amplification can be found at http://ccdb.ca/site/wp-content/uploads/2016/09/CCDB_Amplification.pdf). A second round of PCR amplification added Ion Torrent sequencing adapters and a unique multiplex identifier (MID) sequence to the 5’ end of amplicons for each sample. Amplified COI fragments were then pooled and single-end sequenced on an Ion Torrent PGM sequencing platform (Thermo Fisher, Waltham, MA, USA). The resulting sequence data was automatically de-multiplexed by the PGM Torrent Browser (Thermo Fisher, Waltham, MA, USA). We removed primer and adapter sequences from the de-multiplexed reads, and discarded reads that consisted of only primer or adapter sequence using CUTADAPT v2.3 (Martin 2011). We used DADA2 (Callahan et al. 2016) in R v4.2.2 (R Core Team 2022) to trim and filter reads, estimate error rates for the filtered and trimmed amplicon dataset, combine identical reads into unique sequences, and infer biologically meaningful (as opposed to spurious) amplified sequence variants (ASVs). In the filtering and trimming step, we removed reads containing any unknown basepairs, truncated reads to 200 bp, discarded reads less than 200 bp, truncated reads at the first instance of a quality score less than 2, discarded reads that match the phiX genome, and discarded reads with greater than 2 “expected errors” after truncation (where the number of “expected errors” is calculated from the per-basepair quality scores for each read). In the inference of ASVs, because Ion Torrent sequencing tends to make homopolymer errors and produces higher numbers of indels (relative to Illumina sequencing), we applied a homopolymer gap penalty of -1 and applied a more restrictive threshold for the number of insertions in one sequence relative to another (BAND_SIZE = 32). We then created an ASV abundance table (with the abundance of each ASV for each fish) and removed chimeric sequences using the “consensus” method. We exported the list of ASVs across all samples to a fasta file and used the JAMP pipeline in BOLDigger (Buchner & Leese 2020) to identify the best-fitting taxonomic identification for each ASV.

We used PHYLOSEQ (McMurdie et al. 2013) to create a data object in R that allowed us to associate the abundance of each taxon identified in each fish’s gut with other sample data (i.e. lake, sex, size, and pelvic phenotype). We removed any taxa identified as brook stickleback or bacteria (the latter being more indicative of gut microbiome than diet). Because many taxa were not identified to genus or species, we agglomerated taxa at the rank of family, while also removing taxa that were not identified to the level of family, using the tax_glom function in phyloseq. We analyzed individual-level variation in gut contents by performing an ordination using non-metric multidimensional scaling (NMDS) and Bray-Curtis distances based on family-level taxonomic assignment.

To test the hypothesis that brook stickleback pelvic spine reduction is associated with a difference in diet, we used two analyses. First, to investigate an overall difference in diet, we conducted a PERMANOVA using the adonis2 function in the vegan package (Oksanen et al, 2022) with 10000 permutations to evaluate whether centroids of the Bray-Curtis distances (calculated using phyloseq, as described above) differed among groups. Second, to investigate differences between groups for specific diet items, we used DESeq2 (Love et al. 2014) to conduct Wald tests of the Log_2_ fold-differences in the normalized abundances of each eukaryote family in the diets of brook stickleback (normalized by individual fish) between lakes and between pelvic phenotypes (with p-values corrected for multiple testing with the Benjamini-Hochberg method; Benjamini and Hochberg 1995). Because we had previously found a significant difference in length between fish collected in near-shore “littoral” traps and fish collected in “limnetic” traps suspended >50m from shore (Willerth et al. 2022), which would lead to collinearity in statistical models investigating the effects of fish length and trap location on diet, we looked for correlations among predictor variables using generalised linear models with a Gaussian error distribution (for length – a continuous numerical variable) or a binomial error distribution (for factor variables – pelvic phenotype, sex, and trap location). Given the results of these tests (details below in the results section) showing significant correlation between length and habitat (but between no other predictor variables), we used only one of either habitat or length in our analyses. In the comparison between lakes (for the PERMANOVA and Wald tests), we specified a model that included lake, pelvic morphology, sex, and scaled total length (i.e. scaled to mean = 0 and variance = 1), but not habitat because no fish were captured in “limnetic” traps in Shunda Lake. Because we expect diet might be affected by allometry (Aguirre et al. 2008, Reimchen et al. 2016), we also included interactions with length in all our diet models. Because of concerns with over-fitting given our sample size, we did not include other interaction terms in our diet models. For the Wald tests, we subsequently conducted separate analyses for each lake, wherein we specified models (one per lake) that included pelvic phenotype (spined or pelvic-reduced), sex, scaled total length (or habitat in the case of Muir Lake), and two-way interactions with length.

### Gill rakers

Fish that had been bleached and stained with alizarin red for morphological analyses (Willerth et al. 2022) were used to investigate variation in gill raker morphology associated with different pelvic phenotypes. We removed the first branchial arches from both sides of each fish with forceps and scissors, cleared away all the gill filaments, and photographed the gill rakers in a plastic weighing boat using a Nikon SMZ-75T Greenough-type stereo microscope with a mounted Imaging Source camera and NIS-Elements D Software (©2021 Nikon Corporation).

Following Kaeuffer et al. (2011), we counted the number of gill rakers on the left and right gill arches, and we measured the length of the second through fourth gill raker from the epibranchial-ceratobranchial joint on the ceratobranchial, which were consistently the three longest gill rakers on each arch. We used generalized linear models in R with a gaussian error distribution to model the effects of fish length, sex, year of capture, lake, and pelvic phenotype on variation in gill raker length or number. As described above, we used length as a proxy for habitat to avoid collinearity between the length and habitat variables. We set contrasts among factors using the contr.sum function, and we used the CAR package (Fox and Weisberg 2019) to generate an ANOVA table with type III sums of squares. Given the larger sample size for our gill raker dataset, and the reasonable possibility that the effect of pelvic reduction on gill raker morphology differs between sexes, between lakes, or between years, we included all two-way interactions in our initial models, and then used a reverse-stepwise approach for model selection, removing any interaction terms that were not significant (p > 0.05).

## Results

A list of all taxa identified via metabarcoding in Muir Lake and Shunda Lake is available in the supplementary material (Table S1 and Table S2). At the population level, brook stickleback in Muir Lake and Shunda Lake had taxonomically diverse diets. We identified thirteen Eukaryote phyla among the Amoebozoa, Archaeplastida, Chromalveolata, Fungi, and Metazoa. The Metazoa was by far the most diverse taxon in both lakes where we identified six different phyla, the most abundant and diverse of which were the arthropods. We identified six arthropod classes: Arachnida, Branchiopoda, Copepoda, Insecta, Malacostraca, and Ostracoda. Insecta was by far the most abundant and diverse class (among all phyla) in the diets of brook stickleback in both lakes, with six orders and sixteen families. We focused on family-level classifications because many taxa were not resolved to genus or species. Prior to statistically evaluating differences in diet between lakes and between pelvic phenotypes, we evaluated correlations among predictor variables. Using a type II sums of squares ANOVA (i.e. evaluating a main effect conditional on all other main effects without significant interaction effects), we found a significant effect of habitat (limnetic or littoral) on fish length in Muir Lake (df = 1, Wald’s *X*^2^ = 4.7997, p = 0.028), and a significant effect of fish length on habitat in Muir Lake (df = 1, Wald’s *X*^2^ = 4.8788, p = 0.027). No other predictor variables were correlated.

Large individual-level differences in diet among brook stickleback obscured any obvious group-level differences in diet between lakes or between pelvic morphologies in an ordination plot (Figure 1). A PERMANOVA test of overall differences in diet did, however, reveal that we can reject the null hypothesis that pelvic spine morphs have the same centroid, indicating that diet does differ significantly between pelvic spine morphs (Table 2). Wald tests of the Log_2_ fold-differences in the abundances of each eukaryote family in the diets of brook stickleback between lakes suggested that there were significant differences in the diets of brook stickleback in Muir Lake compared to Shunda Lake attributable to several taxa (Figure 2). Several families found in Muir Lake were not found in Shunda Lake and vice versa: Unionicolidae, Culicidae, Muscidae, Chaoboridae, Hydrozetidae, Ascarididae, Succineidae, Planorbidae, Crambidae, Candonidae, Naididae, Gelechiidae, Chlorellaceae, Scirtidae, and Cyclopidae were not found in Shunda Lake, whereas Simuliidae, Baetidae, Tephritidae, Sellaphoraceae, Saprolegniaceae, Leptolegniaceae, Spionidae, Cyprididae, and Bucculatricidae were not found in Muir Lake. The diet differences between lakes were not driven entirely by the absence of taxa in one lake – several taxa that were present in both lakes (e.g. Caenidae) were consumed in greater abundance in one lake relative to the other (Figure 2).

**Figure 1.**
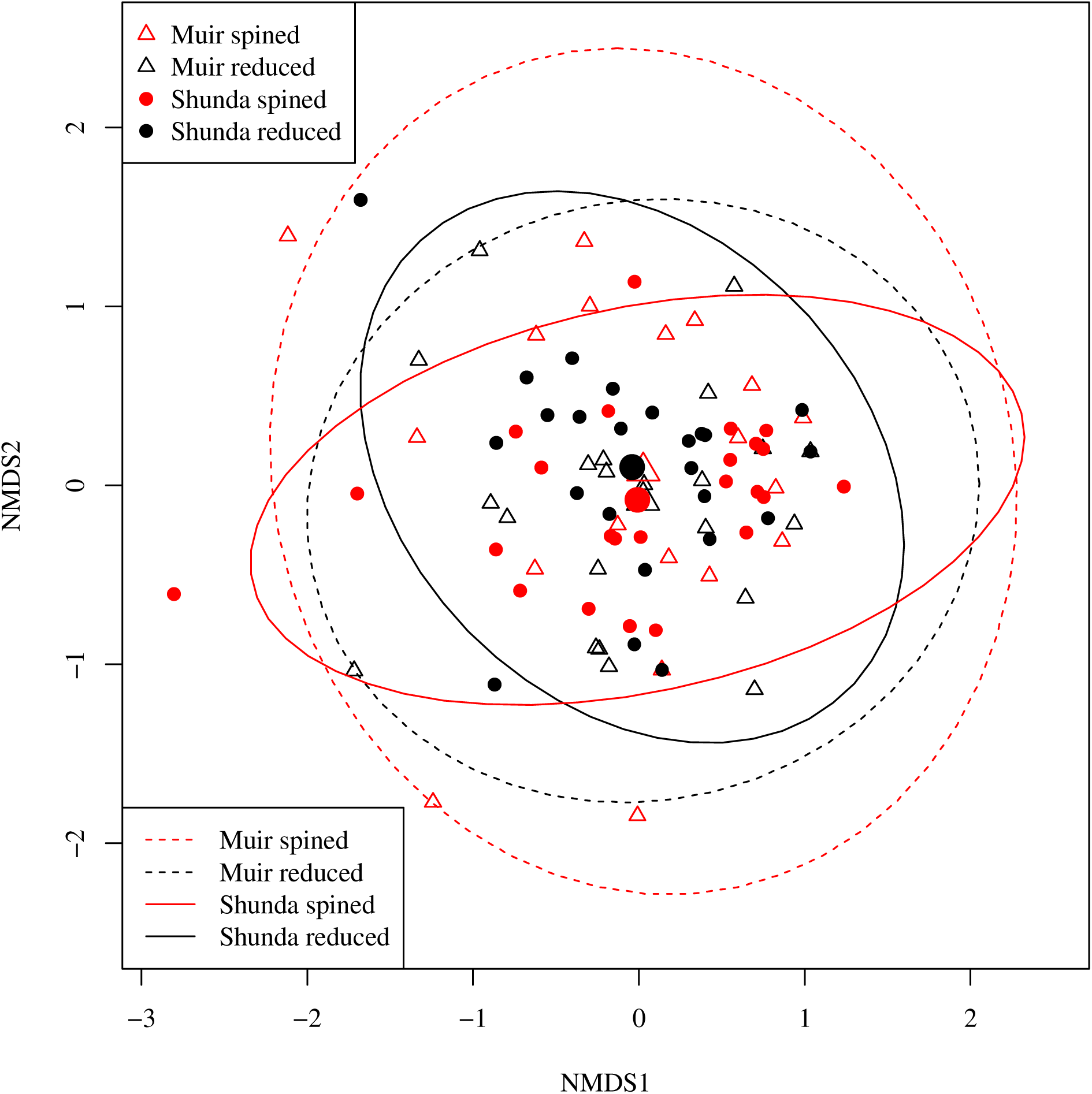
Ordination plot of Non-metric Multidimensional Scaling (NMDS) axes based on sample-wise Bray-Curtis distances in brook stickleback diet (with diet items classified to the level of family). Ellipses show the 95% probability interval for each group, assuming a normal distribution of points. The larger points indicate the center of each ellipse.

**Figure 2.**
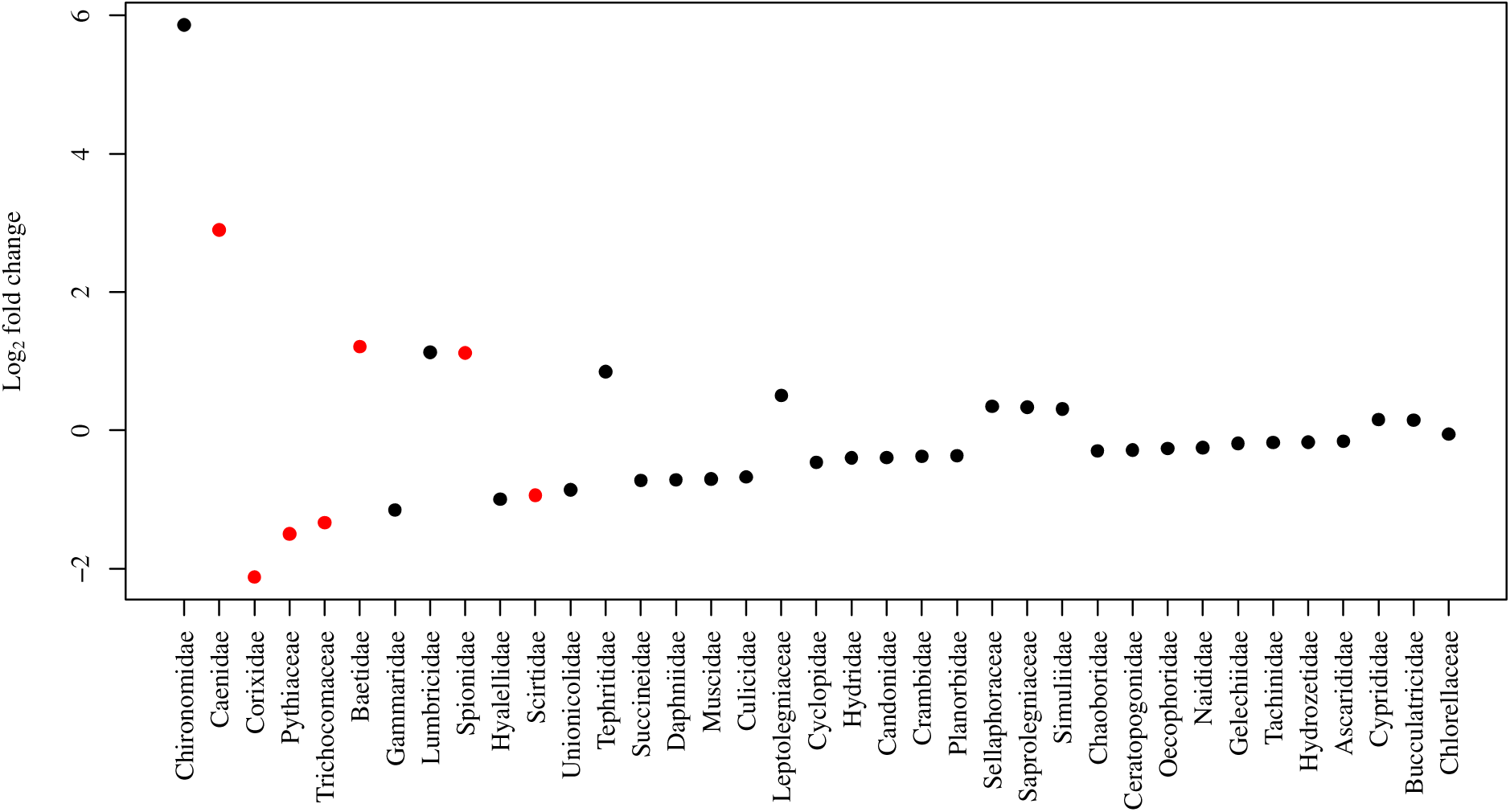
Log_2_ fold change between Shunda Lake and Muir Lake in the abundance of eukaryotic families in the diets of brook stickleback. Zero indicates no difference in abundance between lakes, positive numbers indicate higher abundance in Shunda Lake, and negative numbers indicate higher abundance in Muir Lake. Red symbols indicate a significant difference in abundance with an adjusted p-value < 0.05 corrected for multiple testing. Families are arranged (left to right on the x-axis) in order of highest to lowest fold change (in an absolute sense) between lakes.

**Table 2.** Results of a PERMANOVA test of diet differences.

|  | df | Sum of Squares | F | p-value |
| --- | --- | --- | --- | --- |
| Main effects: |  |  |  |  |
| Lake | 1 | 0.648 | 1.549 | 0.074 |
| Pelvic reduction | 1 | 0.902 | 2.154 | 0.008 |
| Sex | 1 | 0.614 | 1.466 | 0.094 |
| Body length | 1 | 0.923 | 2.205 | 0.008 |
| Interaction effects: |  |  |  |  |
| Lake x Body length | 1 | 0.49 | 1.172 | 0.256 |
| Pelvic reduction x Body length | 1 | 0.649 | 1.550 | 0.075 |
| Sex x Body length | 1 | 0.271 | 0.647 | 0.877 |
| Residual | 86 | 35.998 |  |  |
| Total | 93 | 40.495 |  |  |

To further address the hypothesis that brook stickleback pelvic spine reduction (or loss) is associated with a difference in diet, we evaluated the differential abundance of each taxon identified in the stomachs of spined and pelvic-reduced brook stickleback in Muir Lake and Shunda Lake. When fish length was included as a factor in analyses of diets in Muir Lake, Daphniidae (daphnia, or water fleas) were significantly more abundant in the diets of spined fish, whereas Caenidae (squaregill mayflies), Gammaridae (amphipods commonly known as “scuds”), and Hyalelidae (amphipods) were significantly more abundant in the diets of pelvic-reduced fish (Figure 3). When habitat (instead of length) was included as a factor in analyses of diets in Muir Lake, only Caenidae were significantly more abundant in the diets of pelvic-reduced fish (Figure 4). In Shunda Lake, Lumbricidae (earthworms) were significantly more abundant in the diets of spined fish (Figure 3). Diets also differed significantly based on fish length and between sexes (Figure 3, Figure 4).

**Figure 3.**
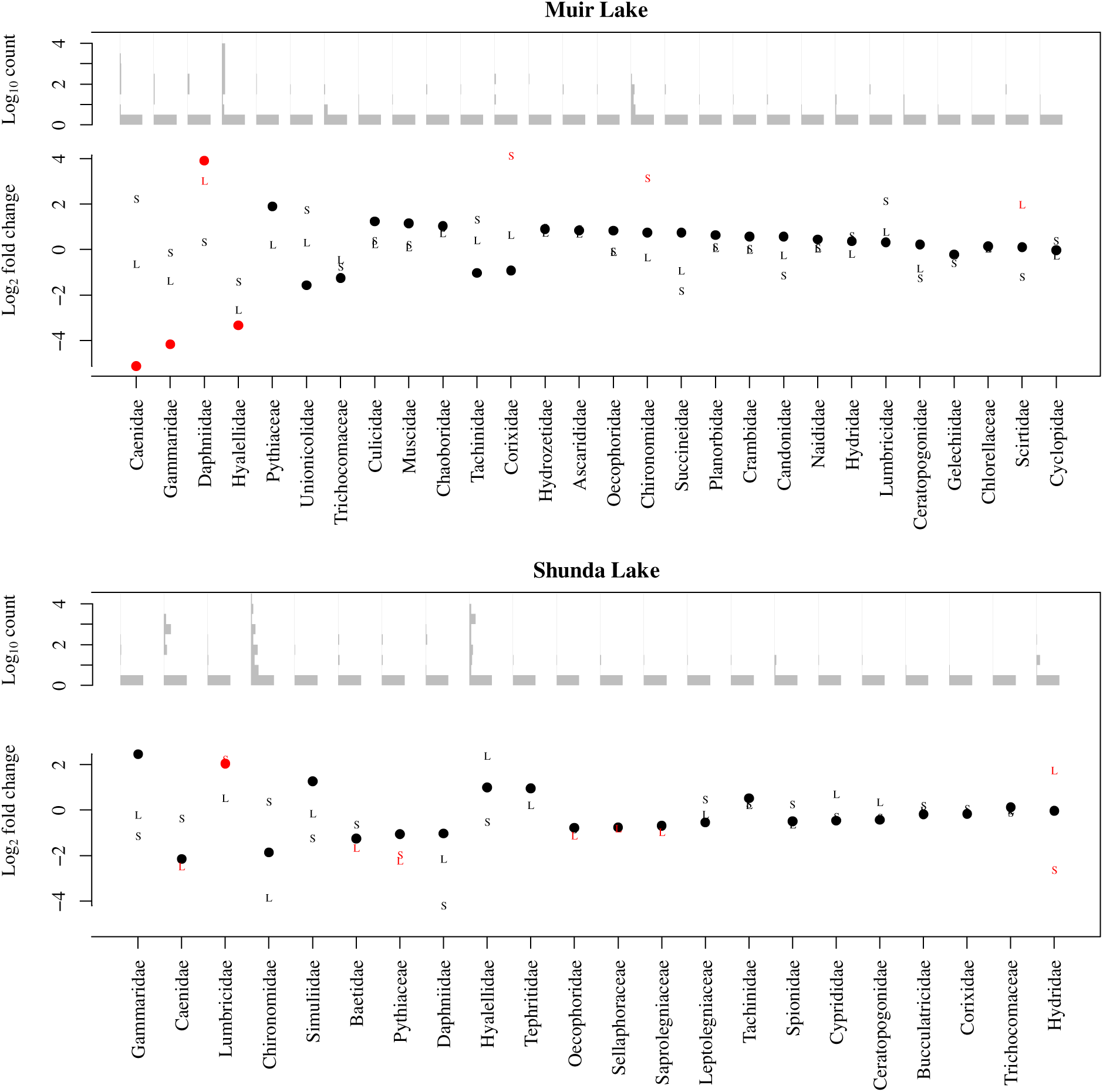
Log_2_ fold change in the abundance of eukaryotic families in the diets of brook stickleback in Muir Lake and Shunda Lake with length as a co-factor. Points show the change in abundance between spined and pelvic-reduced individuals, “S” symbols indicate change between sexes, and “L” symbols indicate change attributed to fish length. A Log_2_ fold change of zero indicates no difference in abundance, whereas positive values indicate higher abundance inpined individuals, in males, or in longer individuals. Red symbols indicate a significant difference in abundance with an adjusted p-value < 0.05 corrected for multiple testing. Two-way interactions with length were included in the statistical model but, for clarity, are not included in this visualization. Families are arranged (left to right on the x-axis) in order of highest to lowest fold change (in an absolute sense) between pelvic morphotypes in each lake. Histograms of Log_10_ abundances across all sampled individuals for each diet item are shown above the fold-change points.

**Figure 4.**
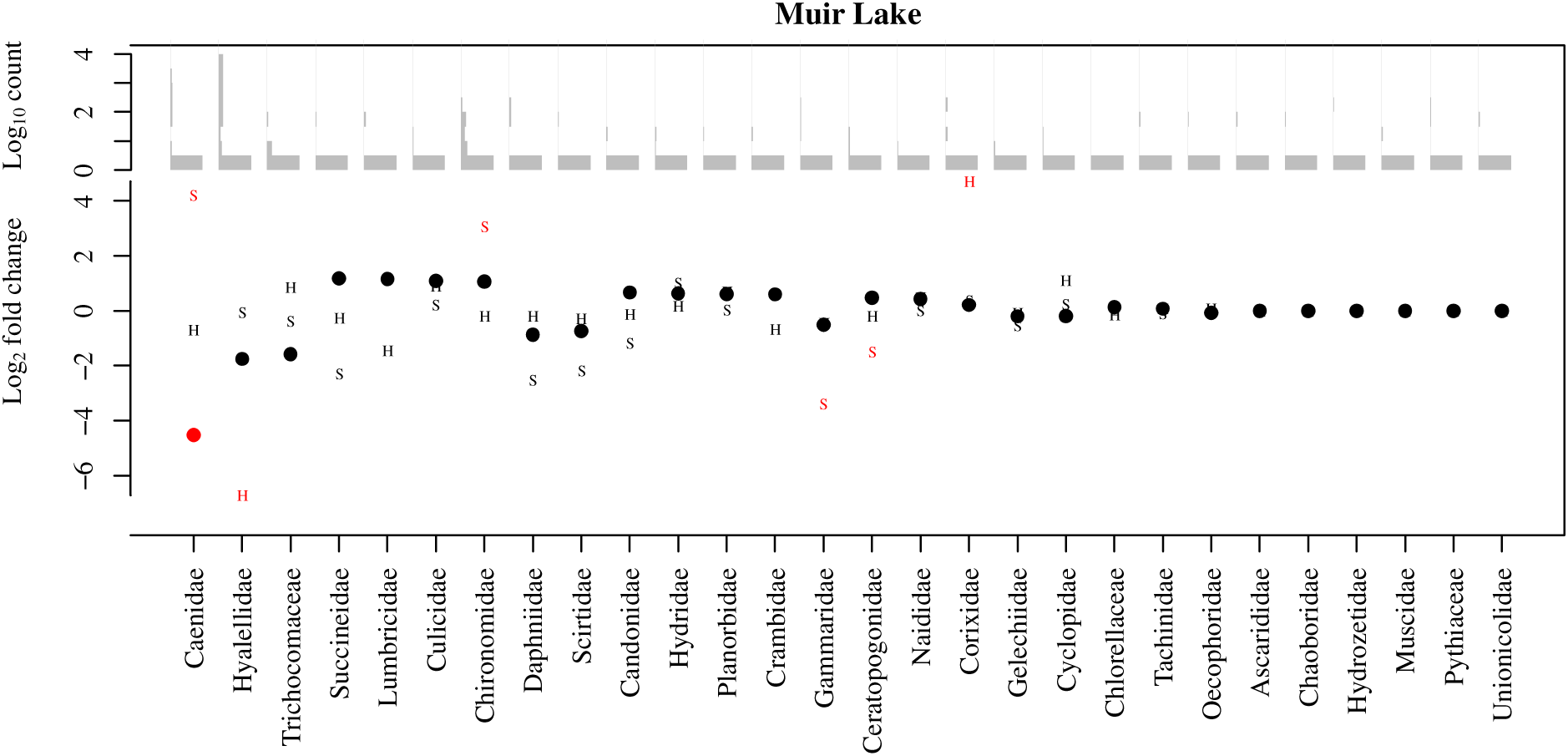
Log_2_ fold change in the abundance of eukaryotic families in the diets of brook stickleback in Muir Lake with habitat as a co-factor. Points show the change in abundance between spined and pelvic-reduced individuals, “S” symbols indicate change between sexes, and “H” symbols indicate change attributed to habitat. A log_2_ fold change of zero indicates no difference in abundance, whereas positive values indicate higher abundance in spined individuals, in males, or in the littoral habitat. Red symbols indicate a significant difference in abundance with an adjusted p-value < 0.05 corrected for multiple testing. Families are arranged (left to right on the x-axis) in order of highest to lowest fold change (in an absolute sense) between pelvic morphotypes in Muir lake. Histograms of Log_10_ abundances across all sampled individuals for each diet item are shown above the fold-change points.

To investigate a potential change in gill raker morphology between spined and unspined brook stickleback, we compared gill raker morphology between individuals with spined and reduced pelvic phenotypes in Muir Lake and Shunda Lake. We conducted separate analyses where we looked for an effect on maximum gill raker length (i.e. the length of the longest gill raker in each individual), mean gill raker length, the length of a specific gill raker (e.g. the second gill raker from the epibranchial-ceratobranchial joint on the ceratobranchial), the number of gill rakers on the left or right gill arch, or the mean number of gill rakers on both arches. We also conducted analyses involving the effect of pelvic morphology with three categories (i.e. spined, vestigial, and unspined) instead of two categories (i.e. spined and pelvic-reduced). Whereas the effects of some factors varied in magnitude among these analyses, no analysis revealed a significant effect of pelvic morphology on gill raker size or number. For clarity, we report only the details of our analyses involving the effect of pelvic reduction (with two categories of pelvic morphology) on maximum gill raker length (Table 3, Figure 5) and on the mean number of gill rakers (Table 4, Figure 6). Gill raker length and number did not differ significantly between spined and pelvic-reduced brook stickleback. Gill raker length and number were significantly affected by fish size, with larger fish having more and longer gill rakers. Brook stickleback in Muir Lake had more gill rakers and longer gill rakers than those in Shunda Lake. Male brook stickleback had significantly longer gill rakers than female brook stickleback, especially at larger sizes. The effect of fish size on gill raker length was weaker for samples collected in 2019 than for samples collected in 2017 (see Tables 3 and 4 for details of statistical results).

**Figure 5.**
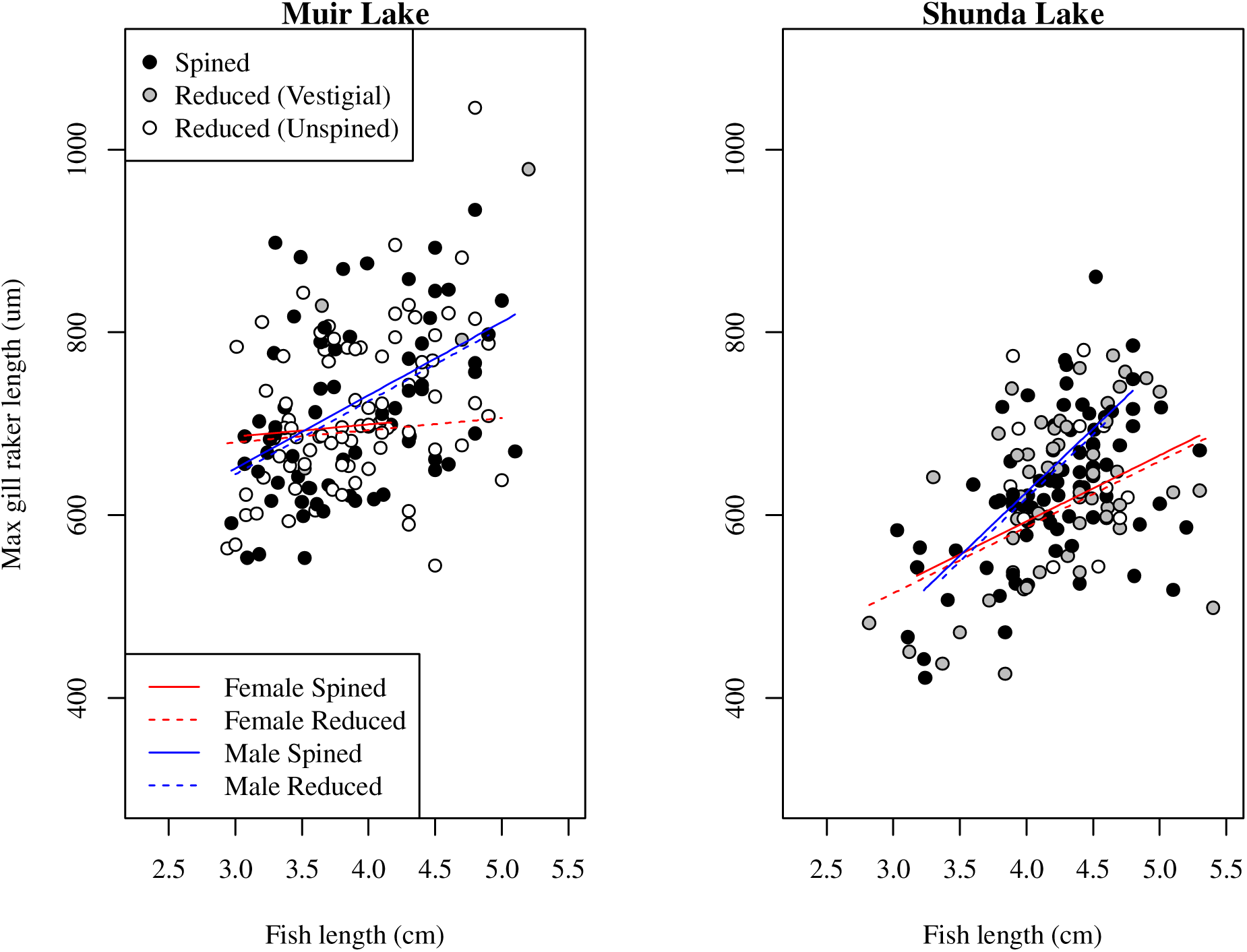
Comparison of maximum gill raker lengths between spined and pelvic-reduced brook stickleback in Muir Lake and Shunda Lake. The horizontal span of each line indicates the range of fish lengths for each group in the comparison (i.e. male spined, female spined, male reduced, and female reduced). For this visualization, only the effects included in the final statistical model (Table 3) are depicted (although, for clarity, the effect of year of sampling is not shown).

**Figure 6.**
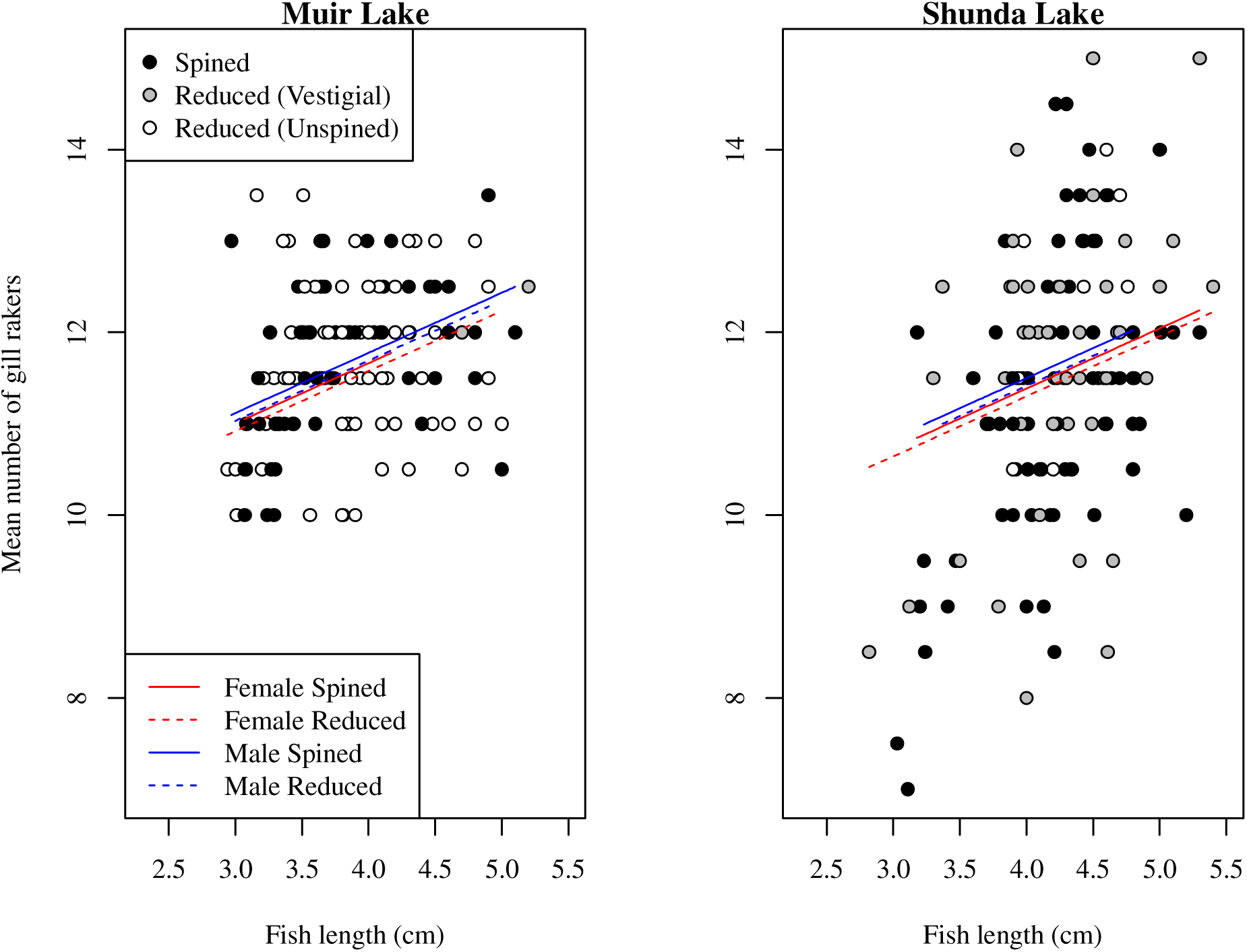
Comparison of mean number of gill rakers between spined and pelvic-reduced brook stickleback in Muir Lake and Shunda Lake. The horizontal span of each line indicates the range of fish lengths for each group in the comparison (i.e. male spined, female spined, male reduced, and female reduced). For this visualization, only the effects included in the final statistical model (Table 4) are depicted (although, for clarity, the effect of year of sampling is not shown).

**Table 3.** ANOVA table for the generalized linear model of the effects of pelvic reduction (spined or pelvic-reduced), lake (Muir Lake or Shunda Lake), sex, sampling year (2017 or 2018), and body length (total length in mm) on maximum gill raker length in brook stickleback. We included all two-way interactions in our initial models, and then used a reverse-stepwise approach for model selection, removing any interaction terms that were not significant. Two-way interaction terms that were not significant (p > 0.05) were not included in the final model shown here.

|  | Sum of Squares | df | *F* | p-value |
| --- | --- | --- | --- | --- |
| Main effects: |  |  |  |  |
| Pelvic reduction | 5109 | 1 | 0.9238 | 0.337 |
| Lake | 112007 | 1 | 20.2544 | < 0.001 |
| Sex | 84370 | 1 | 15.2569 | < 0.001 |
| Year | 16909 | 1 | 3.0578 | 0.081 |
| Body length | 417938 | 1 | 75.5767 | < 0.001 |
| Interaction effects: |  |  |  |  |
| Lake x body length | 55977 | 1 | 10.1224 | 0.002 |
| Sex x body length | 109862 | 1 | 19.8665 | < 0.001 |
| Year x body length | 28198 | 1 | 5.099 | 0.025 |
| Residual | 1548394 | 280 |  |  |
| Total | 2378764 |  |  |  |

**Table 4.** ANOVA table for the generalized linear model of the effects of pelvic reduction (spined or pelvic-reduced), lake (Muir Lake or Shunda Lake), sex, sampling year (2017 or 2018), and body length (total length in mm) on mean number of gill rakers in brook stickleback. We included all two-way interactions in our initial models, and then used a reverse-stepwise approach for model selection, removing any interaction terms that were not significant. Interaction terms were not significant (p > 0.05) and were not included in the final model shown here.

|  | Sum of Squares | df | *F* | p-value |
| --- | --- | --- | --- | --- |
| Pelvic reduction | 0.64 | 1 | 0.5474 | 0.460 |
| Lake | 5.65 | 1 | 4.8253 | 0.029 |
| Sex | 1.52 | 1 | 1.3004 | 0.255 |
| Year | 2.05 | 1 | 1.7528 | 0.187 |
| Body length | 29.06 | 1 | 24.8229 | < 0.001 |
| Residual | 331.28 | 283 |  |  |
| Total | 370.20 |  |  |  |

## Discussion

Whereas the factors maintaining the pelvic spine polymorphism within brook stickleback populations are unclear, it is possible, if not likely, that spined and unspined fish differ ecologically. A previous study of stable isotope signatures revealed that spined and pelvic reduced brook stickleback are associated with different habitat types (Willerth et al. 2022), suggesting that the pelvic spine polymorphism may have ecological correlates. In this study, we tested the hypothesis that spined and pelvic-reduced brook stickleback have different diets. Consistent with our hypothesis, we found significant differences in the abundance of some taxa in the diets of spined and pelvic-reduced brook stickleback.

Pelvic spine reduction in brook stickleback and threespine stickleback populations has evolved in parallel, likely driven by similar selective forces. The extent to which we should expect parallelism to extend beyond a particular phenotype of interest (in this case, pelvic spine reduction), resulting in parallel ecological, behavioural, and morphological consequences, is unknown. We predicted that, like threespine stickleback, pelvic spine reduction in brook stickleback should be associated with a more benthic diet. Our observations did not align with this prediction, which implies non-parallelism in the ecological correlates of spine reduction. Diet differences between spined and pelvic-reduced brook stickleback did not indicate a clear or consistent trend towards a benthic or limnetic diet in either morph. The taxon-specific differences in diet between spined and pelvic-reduced brook stickleback that we report here should not, however, be over-interpreted. The specific diet items that differed significantly between pelvic morphs were highly dependent on choices in our statistical analyses. For example, we note that our choice whether to used fish size or habitat as a variable in our model for Muir Lake influenced our inferences, although this was likely partly due to the reduction in power associated with replacing a continuous variable (length) with a factor variable (habitat). In addition, whether we included more or fewer two-way interactions in our analyses also had a strong effect on which specific taxa differed significantly between pelvic morphs (these alternative analyses are not presented here). Furthermore, our metabarcoding approach did not allow us to differentiate taxa that were actually targeted by stickleback from taxa that happen to be in the guts of the stickleback’s prey. For example, Pythiaceae, known as water moulds, and which we identified in the diets of brook stickleback, are parasites of plants and animals, and may have been found in the guts of mayflies (or other herbivorous prey), or may have been ingested as part of some other diet item, such as rotting vegetation. Our metabarcoding approach also did not allow us to distinguish different prey life stages, which may be associated with different habitats, making it difficult to make inferences regading stickleback foraging habitat based on diet. Nonetheless, all of our analyses indicate some significant difference in diet between spined and pelvic-reduced brook stickleback, providing support to the possibility that pelvic reduction has ecological consequences, such as a difference in foraging habitat.

Differences in diet associated with pelvic spine reduction in threespine stickleback are frequently accompanied by differences in gill raker morphology (McPhail 1984, Ridgeway and McPhail 1984, Schluter 1993, Nagel and Schluter 1998, Rundle et al. 2003, Matthews et al. 2010). Longer, more slender, more numerous gill rakers are better for planktonic feeding, whereas shorter, thicker, less numerous gill rakers are better for benthic feeding. In brook stickleback, we did not observe a significant difference in gill raker morphology between spined and pelvic-reduced fish. There are several possible reasons why the difference in diet between spined and pelvic-reduced brook stickleback is not associated with a difference in gill raker morphology. First, the diet difference associated with pelvic reduction in brook stickleback does not necessarily involve a decrease in planktonic feeding. Second, spined and pelvic-reduced brook stickleback are not reproductively isolated, and each lake likely contains a panmictic population (Lowey et al. 2020). So, in the absence of pleiotropic effects of the spine polymorphism, divergence (or polymorphism) in other traits that are not directly affected by selection is unlikely to evolve. Lastly, the diet differences between spined and pelvic-reduced brook stickleback may not induce any developmental plasticity in gill raker morphology.

It is unclear whether the diet differences between spined and pelvic-reduced brook stickleback are, in fact, associated with a difference in habitat. It is possible that different pelvic morphs use different resources in the same habitat. If this were the case, then maintenance of spine polymorphism must depend on predators in the same habitat exerting balancing selection pressure simultaneously or in a temporally segregated manner, or on a balance between predation pressure and selection based on physiological costs in calcium-deficient environments. A fruitful avenue of future research would likely be an investigation of temporally varying predation on brook stickleback between seasons, between years, and across multiple generations.

Observations of parallel evolution, such the pelvic spine reduction in multiple stickleback species, have long been interpreted as evidence for the involvement of natural selection as a mechanism of evolutionary change. Evolutionary biologists are now trying to understand when, and when not, to expect the evolution of parallel phenotypes to be associated with parallel ecological consequences (e.g. the same behavioural or habitat divergence) and parallel biological pathways (e.g. the same genes or the same mutations). In the case of threespine stickleback pelvic spine reduction, there seems to be parallelism among populations in the genetic basis of spine reduction, and there appears to be parallelism in the ecological consequences and correlates of spine reduction (Miller et al. 2015). Our comparison with instances of pelvic spine reduction in brook stickleback suggests little parallelism between species. Whether, and why, the extent of parallelism might generally be lower in between-species versus between-population comparisons, despite the likelihood of very similar selective scenarios (e.g. based on balancing or divergent selection due to predation in freshwater environments), will undoubtedly be the focus of future research.

## Acknowledgments

We thank two anonymous reviewers and the associate editor who reviewed a previous version of this manuscript for constructive feedback. Samples were collected with the assistance of Alex Farmer, Carolyn Ly, Alyce Straub, Bryce Carson, Moroni Lopez Vasquez, and Kaylee Olszewski. A Natural Sciences and Engineering Research Council of Canada Discovery Grant awarded to J.A.M. supported this work, and E.Y. was supported by an NSERC URSA.

## Data Availability

Private URL for DRYAD archive during peer review (Clicking the link immediately launches a download of the data files): https://datadryad.org/stash/share/CJuLVp4HYMFaQMBDsCUnMMhK_jh4j4V-eSz9Qp6T21E

